# Pilot-scale production of leucine from CO_2_

**DOI:** 10.1101/2025.11.05.686735

**Authors:** Christian Fink, Franziska Steger, Sebastian Schmidt, Elisa Hilts, François Unger, Aquilla Ruddyard, Maximilian Klein, Bart-Jan Akerboom, Thomas Stehrer-Polašek, Stefan Drießler, Shubham Gurav, Ross T. Fennessy, Justin Smith, Günther Bochmann, Simon K.-M. R. Rittmann

**Affiliations:** Arkeon GmbH, Technopark 1, 3430 Tulln a. d. Donau, Austria; ACIB – Austrian Centre of Industrial Biotechnology, Muthgasse 11, 1190 Vienna, Austria; Environmental Biotechnology Group, University of Tübingen, Tübingen, Germany; Archaea Physiology & Biotechnology Group, Department of Functional and Evolutionary Ecology, Universität Wien, Djerassiplatz 1, 1030 Wien, Austria

## Abstract

On a cellular level, proteinogenic amino acids (AAs) are the building blocks of proteins. On a global scale, AAs serve as important nutrients for humans and animals. Beyond their nutritional value, AAs are of relevance in medicine, and, due to their chemical properties, they are indispensable for applications in many different realms, such as pharmaceutics, cosmetics, animal feed, food, and the beverage industry. Here, we report the first archaeal cell factory for leucine production from CO_2_ that has been generated by rational design, random mutagenesis and pathway engineering. The cell factory has been bioprocess-technologically examined and successfully scaled-up with regard to productivity, product quality, and operational stability. In a 2-day fed-batch campaign, we produced 181 g of leucine from CO_2_ at 150 L pilot-plant scale at a mean volumetric leucine productivity of 65 mg L^-1^ h^-1^. A thorough techno-economic analysis indicates that the roll-out of leucine production from CO_2_ is nearly economically feasible on a global scale.

## Introduction

Twenty proteinogenic amino acids (AAs) are relevant for feedstock and human nutrition. Additionally, AAs are of importance in several industrial sectors^1,2^. Especially the essential AAs, including the BCAAs (valine, leucine, and isoleucine), which cannot be synthesized by animals or humans, are of specific interest for industrial AA production. Besides their essential function, BCAAs are valuable in sports nutrition for enhancing muscle growth, reducing fatigue, and accelerating recovery after exercise^3–5^. Additionally, there is evidence for a dietary effect that leucine might be associated with fat reduction^6,7^. The large variety of applications for AAs results in a global market size of USD 32.19 billion in 2025 and is projected to grow to USD 56.61 billion by 2032, with a robust compound annual growth rate (CAGR) of 8.4%^8^. Leucine, the key BCAA relevant for this study, has a market size of USD 1.23 billion in 2025 and is expected to surge to USD 3.67 billion by 2032, reflecting an impressive CAGR of 16.89%^9,10^.

Nowadays, most of the essential AAs, including leucine, are produced in microbial fermentation campaigns^11–14^. The main microbial cell factories for AA production are *Escherichia coli* and *Corynebacterium glutamicum*. Thereof, genetically-engineered strains were optimized over decades for glucose-based production of individual AAs^14–16^. However, due to foreseeable, climate crisis-based restrictions in agricultural land use for glucose-production^17,18^, valorization of alternative carbon sources and C1 compounds, such as hemicellulose, methanol or carbon dioxide (CO_2_) are considered for next generation AA production. To date, approaches for usage of alternative carbon sources, such as xylose or arabinose in existing *E. coli* and *C. glutamicum* cell factories for AA generation have been considered^20,21^. Additionally, C1-fixing microbes as hosts for AA production, such as *Bacillus methanolicus* for lysine production from methanol or *Cupriavidus necator* and *Synechocystis* spp. for valine and aromatic AA production from CO_2_ are employed^22–25^. Recently however, it has been found that CO_2_-fixing methanogenic archaea (methanogens) naturally secrete AAs^26,27^.

One of them is *Methanothermobacter marburgensis*, a thermophilic methanogen, that has already gained attraction as a microbial cell factory for gas-fermentative methane production in Power-to-Gas (P2G) systems^28–30^. In this regard, molecular hydrogen (H_2_) that is generated from renewable energy sources (“green H_2_“), such as wind or solar energy, is used to reduce carbon-captured or waste-gas-derived CO_2_. Hence, H_2_ is biologically catalyzed to methane in P2G or in power-to-chemicals (P2C) to e.g., AAs^31,32^. Within the P2G and in the AA production framework, it has already been proven that bioprocesses with *M. marburgensis* are scalable (Haslinger, Reischl *et al*., submitted for publication). Moreover, *M. marburgensis* is highly resistant to shear forces^28^ due to the rigid characteristics of the archaeal peptidoglycan-harboring cell wall^33^. Recently, markerless mutagenesis tools have been developed that allow metabolic and pathway engineering of *M. marburgensis* to improve production rates of individual AAs^34,35^. Taken together, *M. marburgensis* is a highly suitable chassis for archaeal cell factory development for AA production from CO_2_.

As a first proof-of-concept, we chose the production of the branched-chain amino acid leucine due to its relevance in human sports nutrition and its application as additive in animal feed^36^. On a physiological level, we aimed to increase the carbon flux from CO_2_ to leucine. In a two-step metabolic engineering attempt an archaeal cell factory was developed through rational design and random mutagenesis. On a bioprocess-technological level, the leucine cell factory has been proven in various gas fermentation campaigns regarding bioprocess stability and functionality for leucine production in lab-scale continuous culture and fed-batch at various bioreactors up to 150 L pilot-plant scale. Through a techno-economic model, we put the leucine productivity and leucine concentrations from pilot scale into an economic context to assess the economic feasibility and the putative commercial impact of our technology on a global scale.

## Results

### A combination of genetic engineering and random mutagenesis leads to leucine overproduction in *M. marburgensis*

In the first step, we evaluated if leucine overproduction and secretion by *M. marburgensis* can be achieved by using a random mutagenesis approach to increase the carbon flux to leucine. Through a high-throughput screening of mutagenized mutants, we found significantly higher leucine concentrations with up to an 8-fold and a 15-fold (p<0.01) increase in specific leucine production rate (q_Leu_) and selectivity compared to wild-type *M. marburgensis* (**Supplemental Figure 1,2,3**). All other detected AAs and the strains’ growth behavior were not significantly impacted by the overproduction of leucine (**Supplemental Figure 1,2,3**). Sequencing of three respective overproduction mutants revealed single-nucleotide polymorphisms (SNPs) in each of their isopropylmalate synthase (IPMS)-encoding genes at a position relevant for release of an allosteric leucine inhibition, but no SNPs in the acetolactate synthase (AHAS) encoding genes were detected (**Supplemental Figure 4)** (**Figure 1D**).

**Figure 1.**
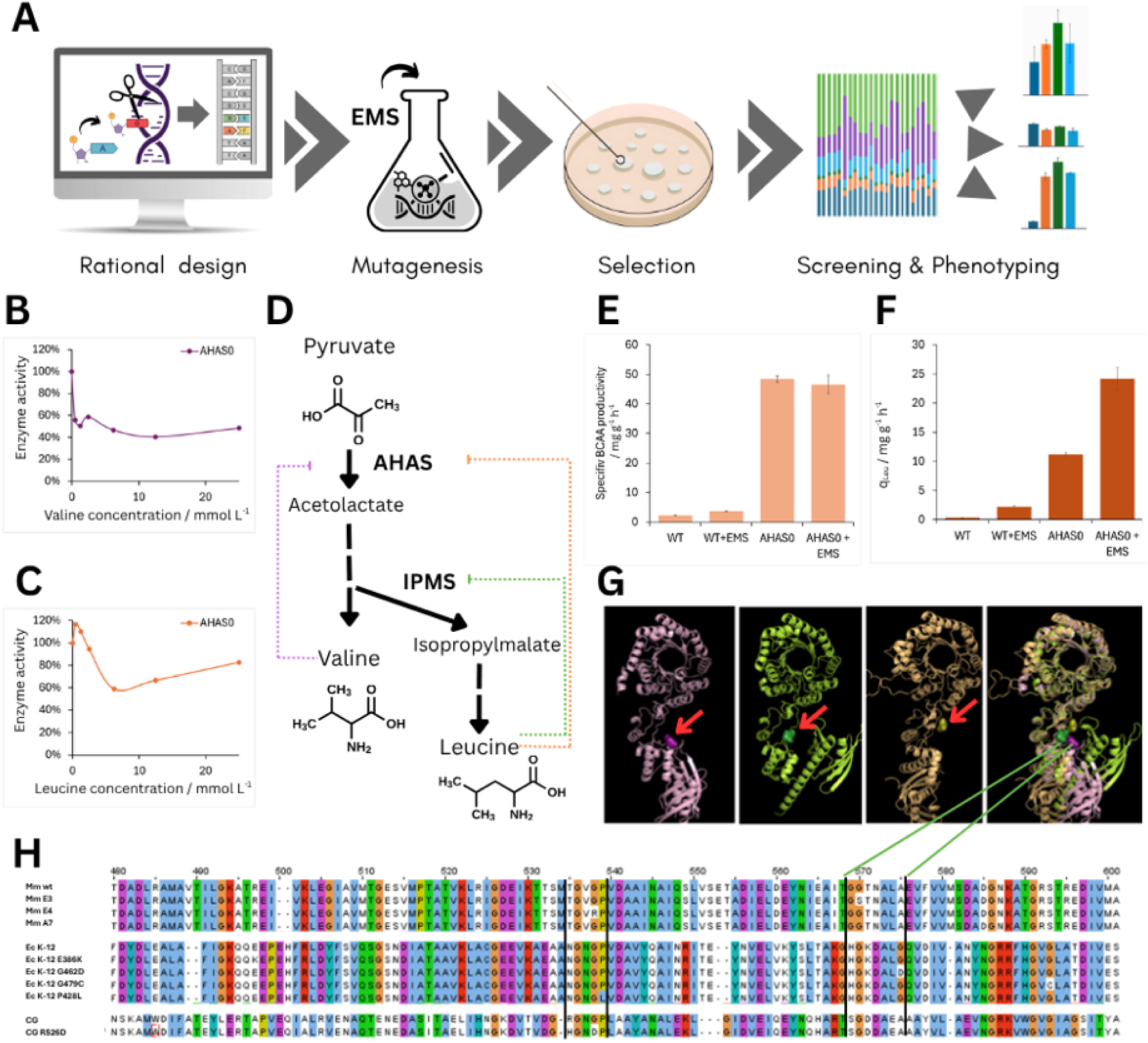
Genetic engineering of M. marburgensis with rational design and random mutagenesis for generation of the leucine cell factory. **A)** Workflow for generation of the leucine cell factory. **B**) Residual activity of heterologous expressed AHAS enzyme with different valine concentrations as allosteric inhibitor measured with in vitro enzyme assay. **C**) Residual activity of heterologous expressed AHAS enzyme with different leucine concentrations as allosteric inhibitor measured with in-vitro enzyme assay. **D**) Schematic valine and leucine biosynthesis pathway from pyruvate. The acetolactate synthase (**AHAS**) and isopropylmalate synthase (**IPMS**) as key allosteric enzymes are depicted as well as the substrate (**Pyruvate**) and the end products (**Valine**, **Leucine**). The dotted lines matching the colors of **B** and **C** represent the allosteric inhibition of the end products on AHAS. The green dotted line matches the allosteric inhibition of leucine on IPMS analyzed in **E** and **G**, **H**) Specific productivity of valine, leucine and isoleucine (**BCAA**) of M. marburgensis wild-type (**WT**), wild-type random mutagenized (**WT**+**EMS**), wild-type with integrated AHAS enzyme from M. thermautotrophicus (**AHAS0**), and the random mutagenized version of AHAS0 (**AHAS0**+**EMS**) in closed batch experiments. The error bars indicate the standard deviation (n=3). **F**) q_Leu_ of the same mutant strains as in **E**. The error bars indicate the standard deviation (n=3). **G**) Cartoon depiction of the IPMS enzymes from M. marburgensis (**pink**), E. coli (**green**), and C. glutamicum (**yellow**) and an overlay of the three. Red arrows indicate the allosteric leucine binding sites analyzed via docking assays with Autodock Vina. **H**) Multisequence alignment of several IPMS enzyme variants. The two areas within black bars indicate the preferred regions for AAs exchanges to result in allosteric inhibition release. The green bars coming from **G** represent the same area as in the predicted leucine binding area.

**Figure 2.**
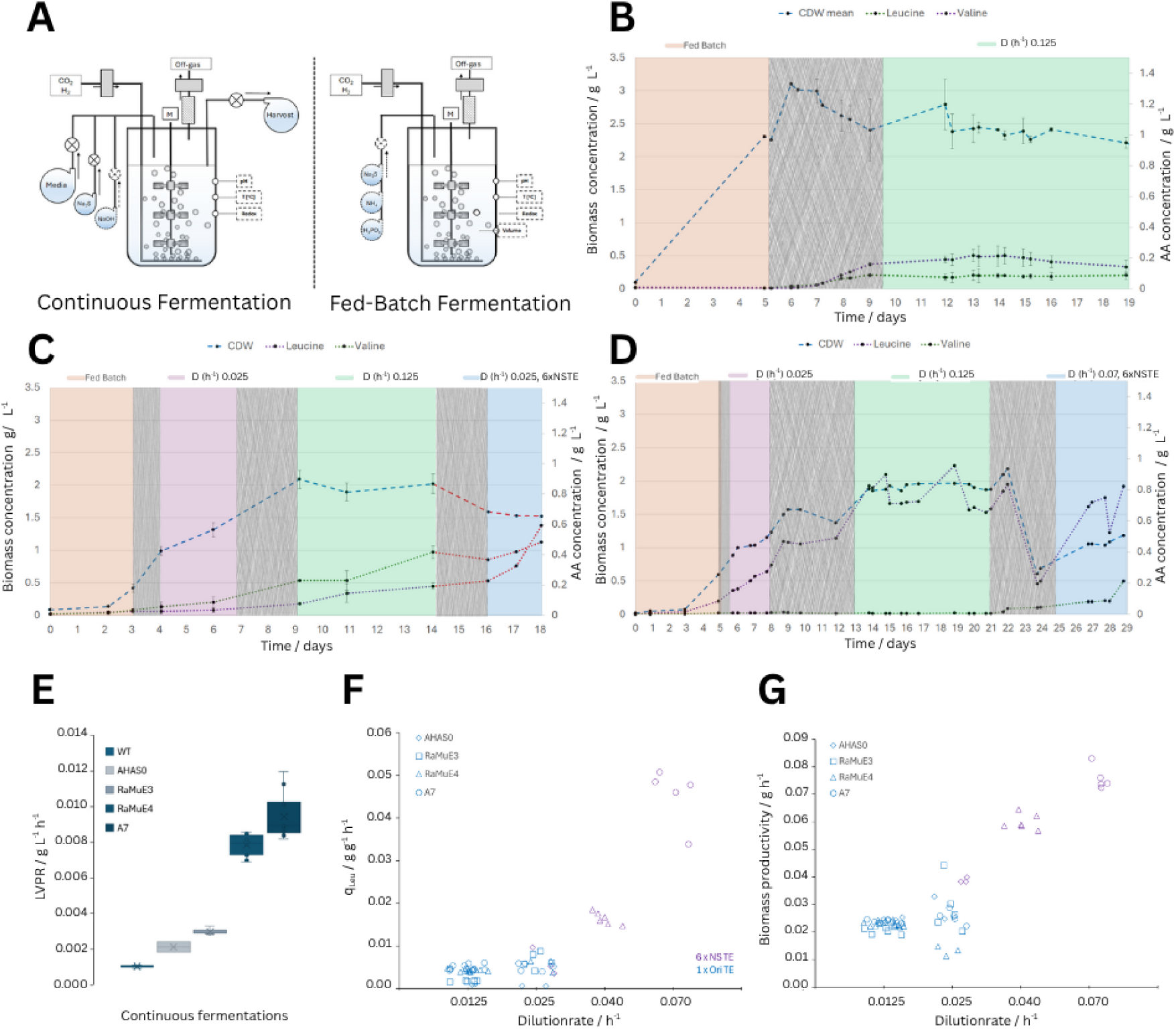
Comparison of stable leucine productivity and growth in continuous culture with wild-type M. marburgensis, M. marburgensis AHAS0, and M. marburgensis AHAS0+EMS mutants and different cultivation conditions. **A**) Schematic depiction of the continuous cultivation of M. marburgensis (**left**) in comparison to the fed-batch cultivation (**right**) described in Figure 3. **B**) Time course depiction of continuous cultivation with M. marburgensis wild-type with an initial fed-batch phase (**orange**), a transition phase (**grey**), and steady state production phase with OriTE as trace elements and dilution rate of 0.0125 h^-1^ (**green**). Error bars mark the standard deviation (n=2). **C**) Time course depiction of continuous culture with M. marburgensis AHAS0 with initial fed-batch phase (**orange**), followed by three different steady state conditions with D of 0.025 h^-1^+OriTE (**purple**), 0.0125 h^-1^+OriTE (**green**), and 0.025 h^-1^+6·NSTE (**blue**). Error bars mark the standard deviation (n=2). **D**) Time course depiction of continuous cultivation with M. marburgensis AHAS0+EMS (**A7**) with initial fed-batch phase, followed by three different steady state conditions with D of 0.025 h^-1^+OriTE (**purple**), 0.0125 h^-1^+OriTE (**green**), and 0.025 h^-1^+ 6·NSTE (**blue**) (n=1). **E**) Averaged LVPR across the different cultivation conditions with M. marburgensis wild-type (**WT**), AHAS0, AHAS0+EMS mutant E3 (**RaMuE3**), AHAS0+EMS mutant E4 (**RaMuE4**), AHAS0+EMS mutant A7 (**A7**). **F**) q_Leu_ across the tested strains and conditions at different D and trace element compositions with M. marburgensis mutants AHAS0, AHAS0+EMS mutant E3 (**RaMuE3**), AHAS0+EMS mutant E4 (**RaMuE4**), AHAS0+EMS mutant A7 (**A7**). **G**) Biomass productivity across the tested strains and conditions at different dilution rates and trace element compositions with M. marburgensis mutants AHAS0, AHAS0+EMS mutant E3 (**RaMuE3**), AHAS0+EMS mutant E4 (**RaMuE4**), AHAS0+EMS mutant A7 (**A7**).

After the first proof-of-concept for overproduction of leucine, we tested whether further increased carbon flux to valine and leucine is achievable by rational design, specifically by genetic engineering via overexpression of the native AHAS0 from *Methanothermobacter thermautotrophicus* in *M. marburgensis* (**Figure 1D**). Here, we demonstrated significantly increased specific valine production rates (q_Val_) and q_Leu_ of 29- and 31-fold (p<0.01, n=3), respectively with the *M. marburgensis* AHAS0 mutant compared to the wild-type strain (**Figure 1E+F**). To analyze the impact of allosteric leucine or valine inhibition on the heterologous AHAS0, we performed an in vitro enzyme assay and found 50% and 35% of inhibition, respectively (**Figure 1B+C**). Thus, although AHAS0 was partially inhibited, these findings render the enzyme still reasonably functional at high leucine and valine concentrations.

To combine the effects of random mutagenesis and overexpression of AHAS0 to enhance leucine production, we performed random mutagenesis on *M. marburgensis* AHAS0 (**Figure 1A**). After screening 36 mutants, we characterized the three most promising leucine-production mutants based on leucine specificity and concentration in closed batch experiments (n=3) (**Supplemental Figure 5,6,7**). We found a 2-3-fold increase in q_Leu_ and a decrease in q_Val_ of up to 5-fold (**Figure 1E+F)** compared to *M. marburgensis* AHAS0. That marks a selectivity towards leucine compared to all other 20 AAs of up to 67% (**Supplemental Figure 5,6,7).** Interestingly, we found SNPs leading to G458S, G426R, and A461V in the *ipms* gene of the randomly mutagenized *M. marburgensis* AHAS0 mutants E3, E4, and A7, respectively. These SNPs occurred in similar locations as they had been already confirmed in allosteric released IPMS from *E. coli* or *C. glutamicum* (**Figure 1H**)^37–39^. To support our findings, we confirmed allosteric leucine binding sites in IPMS with docking assays using Autodock Vina. Although the regulatory domain of IPMS is less conserved across the bacterial and archaeal domains compared to the catalytic domain of IPMS, the allosteric binding site of leucine appears to be highly similar (**Figure 1G**).

Hence, the randomly mutagenized *M. marburgensis* AHAS0 mutants E3, E4, and A7 were considered the first archaeal leucine cell factory and transferred bioreactor systems for continuous cultivation.

### Continuous gas fermentation with *M. marburgensis* leucine cell factories result in stable leucine overproduction with up to 80% leucine specificity

In closed batch cultivations we proved that randomly mutagenized *M. marburgensis* AHAS0 mutants had increased q_Leu_ with high specificity for leucine (**Supplemental Figure 5**). However, relevant insights for industrial processes can only be generated through bioreactor-based cultivation. We compared a duplicate of wild-type *M. marburgensis* (**Figure 2B**) to a duplicate of *M. marburgensis* AHAS0 and the isolated random mutagenized mutants E3, E4, and A7. We applied different liquid dilution rates (D) of 0.0125 h^-1^, 0.025 h^-1^, 0.04 h^-1^, or 0.07 h^-1^ and trace-metal compositions to find suitable cultivation conditions for high volumetric- and/or specific productivity as well as high leucine specificity. Besides the validation of increased productivities, we aimed to identify which of the mutagenized *M. marburgensis* AHAS0 strain performs best for H_2_/CO_2_ conversion to leucine.

With *M. marburgensis* AHAS0 and the A7 mutant, we tested three different cultivation conditions (**Figure 2C+D**). We found that the q_Leu_ of A7 in closed batch experiments suitably matches the results from continuous cultivations with 0.025 g g^-1^ h^-1^ and 0.043 g g^-1^ h^-1^, respectively (**Figure 1F+G; Table 1**). Additionally, the observed fold-increases in leucine concentrations in A7 compared to the wild-type *M. marburgensis* (20-fold) and the AHAS0 mutant (1.7-fold), were highly similar to the results from closed batch experiments. The volumetric leucine production rate (LVPR) of E4 and especially of A7 of up to 0.012 g L^-1^ h^-1^ exceeded the wild-type *M. marburgensis* volumetric productivity by 10-fold and the native AHAS mutant by 4-fold on average (**Figure 2E**). Additionally, we demonstrated a positive influence of a high D (>0.04 h^-1^) in combination with 6·NSTE trace element composition on q_Leu_ with the highest measured productivity in A7 of 0.05 g g^-1^ h^-1^ (**Figure 2F**). The biomass productivity follows the same pattern as q_Leu_ (**Figure 2G**). However, while we experienced a linear correlation in the biomass productivity, we identified an exponential correlation for the q_Leu_ (**Figure 2F+G**). From our results with continuous cultivation, we concluded that mutant A7 provides highly stable leucine production over a period of at least 8 d (*e.g*., at day 13-21 (**Figure 2D**)) and the highest q_Leu_ and LVPR. Therefore, we chose A7 as the leucine cell factory for further investigation.

**Table 1.** Comparison of key concentrations, productivity, and leucine percentage in continuous- and fed-batch fermentations at different scales with the leucine cell factory of M. marburgensis strain A7.

| Gas fermentation | $x / \text{g L}^{-1}$ | LVPR / $\text{g L}^{-1} \text{ h}^{-1}$ | $q_{\text{Leu}} / \text{g g}^{-1} \text{ h}^{-1}$ | Leu / % | Leu / $\text{g L}^{-1}$ | LVPR $\text{MER}^{-1}$ |
| --- | --- | --- | --- | --- | --- | --- |
| Continuous 1.6 L (n=1) | 1.19 | 0.049 | 0.044 | 68.33 | 0.82 | n/a |
| Fed-Batch 1.6 L (n=3) | 3.29 | 0.052 | 0.019 | 67.56 | 1.80 | 0.0045 |
| Fed-Batch 10 L (n=1) | 3.39 | 0.043 | 0.014 | 65.12 | 1.91 | 0.0037 |
| Fed-batch 100 L (n=1) | 4.88 | 0.065 | 0.017 | 76.66 | 1.81 | 0.0057 |

### High leucine concentrations are obtained in fed-batch

In continuous culture we were able to prove consistent q_Leu_ and LVPR with *M. marburgensis* AHAS0 mutant A7. In addition, we have shown that leucine production is stable. However, continuous cultivation is preceded by an initial fed-batch phase and, after a continuous culture is initiated, it takes 3-volume-exchanges to reach steady-state-like conditions. Furthermore, in continuous culture lower total concentrations of products are reached due to the continuous outflow of reactor broth. To prove increased leucine concentrations alongside a shortened bioprocess operation, we set up a fed-batch using 6·NSTE as trace-metal composition. The main differences between the continuous culture and the fed-batch process were the continuous feed and effluent of media combined with continuous supply of sodium sulfide and pH regulating sodium hydroxide, which had been adapted to the biomass concentration throughout the fed-batch process (**Figure 2A**). We performed fed-batch cultivation with A7 in 1.6 L-scale (n=3) (**Figure 3C**). Here, we reached an average LVPR of 0.052 g L^-1^ h^-1^, demonstrating the same volumetric productivity as with 6·NSTE at a dilution rate of 0.07 h^-1^ of 0.049 g L^-1^ h^-1^ in continuous culture (**Table 1**). The final leucine concentration was up to 1.8 g L^-1^ which marks a 2.25-fold increase compared to maximum leucine concentration in continuous culture. However, we obtained the same leucine specificity of 68% in fed-batch and in continuous culture (**Table 1**). While a similar volumetric productivity was observed, the lower biomass of 1.19 g L^-1^ in continuous culture compared to 3.29 g L^-1^ in fed-batch cultivation lowered q_Leu_ in fed-batch fermentation (0.019 g g^-1^ h^-1^) compared to continuous fermentation (0.044 g g^-1^ h^-1^) (**Figure 2D**; **Figure 3C**; **Table 1).**

**Figure 3.**
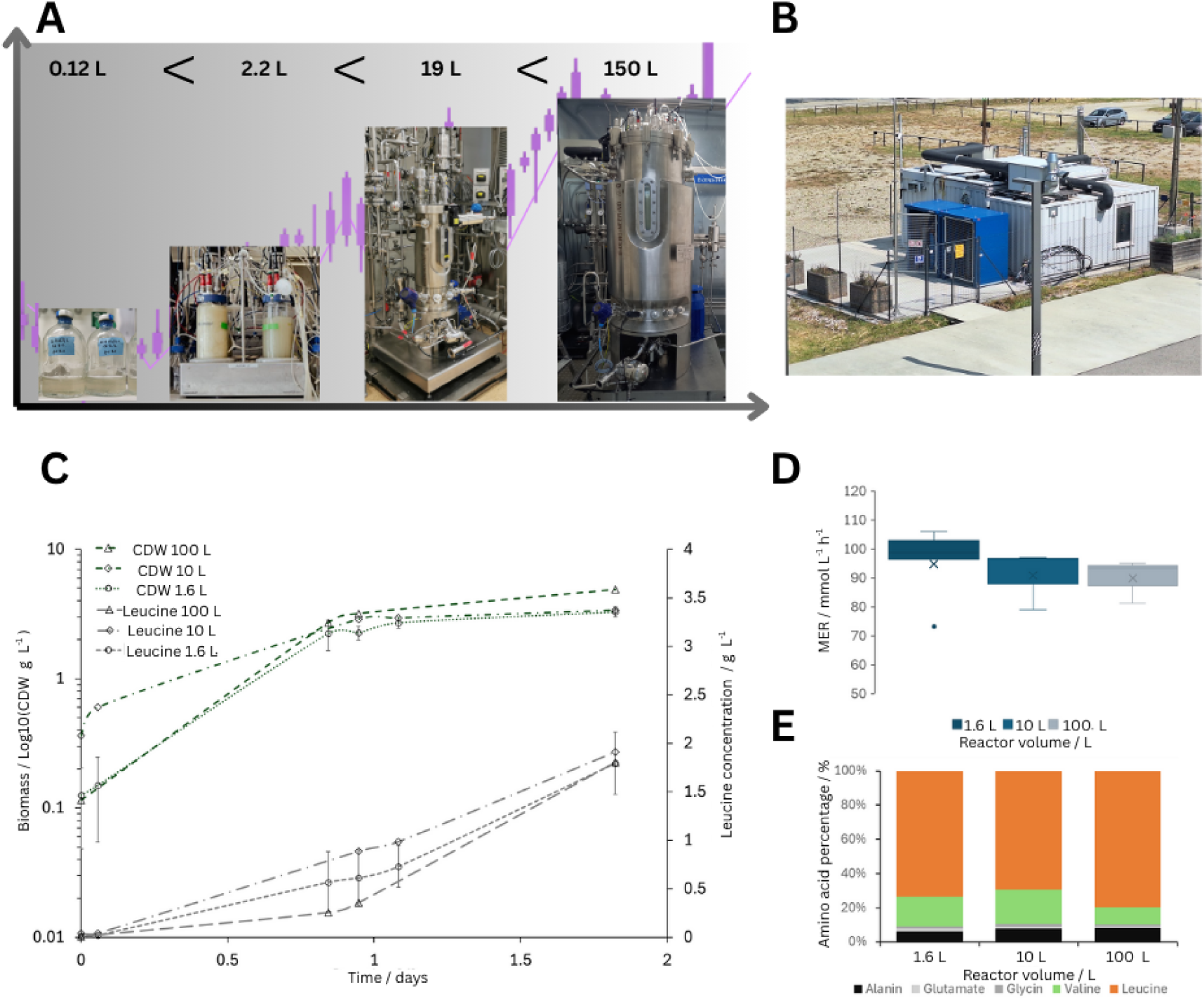
Scalability of the fed-batch process regarding growth and leucine production with the leucine cell factory of M. marburgensis. **A**) Different scales tested from 0.12 L (0.02 L working volume), 2.2 L (1.6 L), 19 L (10 L), to 150 L (100 L) reactor volume, with pictures of the respective serum bottles or reactor vessels (and their vessel volume). **B**) Arkeon GmbH pilot plant in Vienna, Austria, with 150 L gas fermentation bioreactor, media preparation room, and downstream processing (DSP) sample preparation. **C**) Time course of biomass concentrations (**gCDW**) and leucine concentrations (**g L^-1^**) during fed-batch cultivations in 1.6 L (**green**), 10 L (**orange**), and 100 L (**purple**). Error bars mark standard deviation in 1.6 L cultivations (n=3). **D**) Averaged MER at different scales from 1.6 L, 10 L, to 100 L. The error bars mark the standard deviation. **E**) Amino acid percentage of relevant AAs, alanine (**black**), glutamate (**light grey**), glycine (**dark grey**), valine (**green**), leucine (**orange**) (above 1% of total AAs) at the end of the fed-batch cultivation in different cultivation volumes of 1.6 L, 10 L and 100 L.

### Leucine productivity and concentrations in fed-batch are scalable up to pilot plant scale

After the validation and verification of our results at lab-scale, we aimed for scale-up of leucine production with A7 to a 100 L working volume in a 150 L bioreactor. Therefore, we scaled up leucine production in fed-batch two steps from 1.6 L to 10 L. Subsequently, after validation of the process in 10 L, we scaled-up to 100 L (**Figure 3A**). In total, a scale-up factor of 62.5 by volume was achieved. To be able to operate on a 100 L scale, we designed and built our own container laboratory in Vienna, *Seestadt Technologiezentrum 2*. Besides the 150 L bioreactor, the container lab is also equipped with media preparation opportunities, sterilization in place (SIP) and cleaning in place (CIP), and basic DSP (**Figure 3B**).

In terms of A7 growth behavior and leucine concentration, we observed highly similar results at all three scales, 1.6 L, 10 L and 100 L (**Figure 3C**; **Table 1**). The similarity between the scales is also visible in terms of methane evolution rate (MER) of 90-105 mmol L^-1^ h^-1^ (**Figure 3D**) and the specificity in leucine production with roughly 70% at all scales (**Figure 3E**, **Table 1)**. Scalability was also achieved in terms of specific productivity, volumetric productivity, and the ratio of MER and leucine production on a C-molar basis (**Table 1**). This resulted in 181 g of leucine produced from CO_2_ within 2 days of bioprocess runtime with the *M. marburgensis* mutant A7 in a single fermentation campaign.

### Techno-economic analyses reveal H_2_ price and the fate of produced methane as the main cost factors of CO_2_-based AA production

To put the pilot-scale production of leucine from CO_2_ in an economic context, we generated a techno-economic model (TEM) allowing us to estimate the cost of production. Within the TEM, two configurations were considered: I) Biomethane generated during the fermentation process was recirculated through a steam methane reformer (SMR). This configuration enables the reconversion of methane into H_2_ and CO_2_, which can subsequently be reintegrated into the gas fermentation process. II) Biomethane was assumed to be sold externally at a market price of 70 € MWh⁻¹^40^. Although this approach generates additional revenue streams, the reduced internal H_2_ availability may increase the overall production costs.

Based on these initial assumptions of the TEM, we conducted sensitivity analyses for different modes of operation with the current fed-batch process setting. In those, we estimated base values for a 1500 m^3^ bioreactor system and 9 crucial process parameters, such as biomass selling price, methane selling price, H_2_ price (green H_2_, grey H_2_), downstream processing (DSP) efficiency, CO_2_ price, biomass production rate (BPR) to MER ratio, LVPR to MER ratio, bioreactor runtime per year, and LVPR (**Figure 4A**). The base value for LVPR of 0.45 g L^-1^ h^-1^ was estimated to be 7-fold higher compared to the measured 0.065 g L^-1^ h^-1^ in the pilot plant, as this reflects our projected performance enhancement through cell factory and process development and scaling effects (**Table 1**).

**Figure 4.**
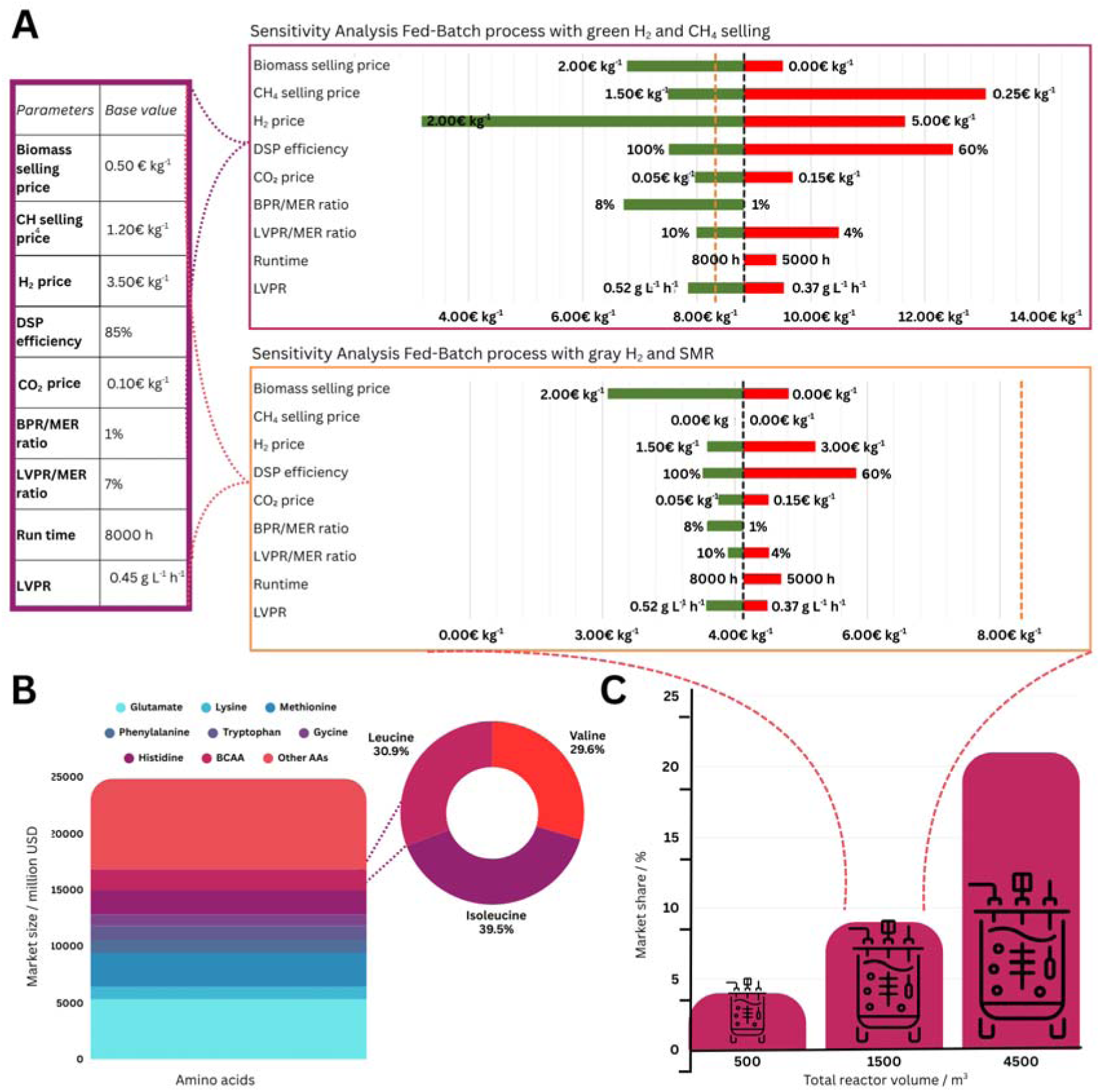
Techno-economic sensitivity analysis of leucine production from CO_2_ with M. marburgensis (**A+C**) and market survey on global amino acid production (**B**). **A**) Techno-economic sensitivity analysis for fed-batch processes generated with base values for biomass selling price, methane (CH_4_) selling price, H_2_ price, DSP efficiency, CO_2_ price, biomass production rate (BPR), and MER ratio, leucine volumetric productivity (LVPR) and MER ratio, the annual bioreactor run time, and the LVPR (**purple box**). A Sustainable scenario with green H_2_ and CH_4_ valorization (**upper right**) and a cost-effective scenario where grey H_2_ and SMR were used (**lower right**) were calculated. The black dashed line marks the production cost of leucine with base values, and the orange dashed line indicates the estimated threshold for economically competitive production cost. Green and red bars show the influence of each individual deviation from base values. **B**) Total market volume in USD for all 20 proteinogenic amino acids with percentual distribution for BCAAs (**donut diagram**). **C**) Estimated market share of leucine produced from CO_2_ with corresponding total bioreactor capacity. The orange dashed lines mark the reactor capacity, on which the sensitivity analyses are based.

In the most sustainable scenario with green H_2_ and selling biomethane, we calculated leucine production costs of 8.80 € kg^-1^ with costs majorly impacted by the price of green H_2_, the DSP efficiency, and the biomethane selling price (**Figure 4A**). Additionally, we estimated the most cost-effective way of leucine production with grey H_2_ and SMR implementation. Here we found a base value cost of production of 4.10 € kg^-1^. In this scenario, the cost was mainly influenced by DSP efficiency, H_2_ price, and the biomass selling price (**Figure 4A**). Taking 8.40 € kg^-1^ as a realistic benchmark for economically feasible production of leucine from our internal market survey (**Figure 4**), the scenario using grey H_2_, and a SMR for energy carrier recycling demonstrates a cost-competitive opportunity for leucine production from CO_2_. The outcome of the sustainable scenario underscores the strong sensitivity of the process to H_2_ pricing. Similarly, the influence of DSP efficiency highlights the importance of product recovery yields in determining the overall process economics. With our currently measured LVPR of 0.067 g L^-1^ h^-1^, 0.6% LVPR MER^-1^, 0.1 mol L^-1^ h^-1^ MER (**Table 1**, **Figure 3**), and 85% DSP efficiency, assuming 2.00 € kg^-1^ H_2_ price, 1.00 € kg^-1^ selling price of biomethane, 0.50 € kg^-1^ biomass selling price, and 1000 m^3^ bioreactor capacity, the production cost of leucine to break even would be 9.35 € kg^-1^.

For a broader context and potential go-to-market strategy in the field of AA production, we conducted a market survey. We found that the production of BCAAs represents about 7.6% of the total AA market size of ∼25 billion USD of around 1.8 billion USD. Leucine contributes 30.9%, valine 29.6% valine, and isoleucine 39.9% (**Figure 4B**). Dependent on the bioreactor capacities of 500 m^3^, 1500 m^3^, or 4500 m^3^, the leucine production process from CO_2_ could contribute 4%, 9%, or even 21% of the total market share, respectively (**Figure 4C**).

## Discussion

In this study, we generated an archaeal cell factory for leucine production from CO_2_ and scaled the production process to a 150 L pilot plant scale. We showed that 2 g L^-1^ leucine was consistently produced across all bioreactor scales with comparable LVPR of 0.05 g L^-1^ h^-1^. Hereby, leucine was produced with high specificity of up to 80% share of AA. With our TEM, we calculated a cost for leucine production of 9.35 € kg^-1^ with 1000 m^3^ reactor capacities, which is close to meeting the internal threshold for economic feasibility of the process of 8.40 € kg^-1^. Our findings unambiguously prove that *M. marburgensis* is a suitable platform for sustainable commodity chemical production from CO_2_.

By the integration of a native AHAS-encoding gene from *M. thermautotrophicus*, we achieved 100-fold increased carbon flux from CO_2_ to the BCAA biosynthesis pathway (**Figure 1F**) even with the responsible AHAS enzyme being product-inhibited by up to 50%, similar to AHAS in *Methanococcus maripaludis*^41^. It has been shown that the release of this product inhibition and fine-tuning of enzyme expression in the leucine biosynthesis pathway have positive effects on the carbon flux from pyruvate to valine production in *E. coli* and *C. glutamicum*^11,42–44^. Neither optimization has been applied to the cell factory which shows strong potential for further development. This can be aided by the recently established genome-scale metabolic model (GEM) for *M. marburgensis*^45,46^. Nevertheless, the presented leucine concentrations and volumetric productivity are by far the highest for biomass derived chemicals produced with archaea, up to date^32,46^ (**Figure 3**, **Table 1**).

The proof of stable productivity in continuous culture, accompanied by similar leucine concentration and volumetric productivity in fed-batch of up to 150 L pilot scale, strongly indicates high genome stability and great potential for large-scale industrial chemical production with genetically engineered *M. marburgensis*. In continuous gas fermentations, we have experienced higher q_Leu_ with less biomass than in fed-batch fermentations (**Table 1**) which indicates gas-liquid mass transfer limitations of H_2_ or CO_2_. Those can be overcome with further process development, such as CFD analysis-aided methods for increased gas availability (Haslinger, Reischl *et al.*, submitted), a complementary method assisted by hybrid models for the GEM-based analysis^45^, or customized reactor designs^47^. For methane production, *Methanothermobacter* spp. have already proven their industrial potential with Electrochaea GmbH or Krajete GmbH^28,32^.

Gaseous methane as the major by-product facilitates DSP of soluble chemicals when using methanogens as production host. One caveat, however, is the high carbon flux to methane compared to leucine (**Table 1**). Hence, as the TEM suggests, methane production needs to be reduced, or the leucine production increased for commercial purposes. A technical solution is the integration of biomethane recirculation via SMR that not only enhances carbon efficiency but also significantly reduces production costs, suggesting that circular gas utilization could represent a key strategy for achieving cost-effective and sustainable AA production in gas fermentation systems. Overall, the sensitivity analysis of the best-case scenario yielded 4.10 € kg⁻¹ of leucine production costs with base values (**Figure 4A**). These results clearly demonstrate that H_2_ pricing and process efficiency, particularly within the SMR and DSP stages, are the dominant cost determinants in the system. This observation aligns with previous techno-economic studies on gas fermentation processes, which similarly identified H_2_ supply and DSP recovery as key economic bottlenecks^48,49^.

The sensitivity analyses based on the TEM support the estimations about green H_2_ and grey H_2_ but also indicate that with higher LVPR and reduced CO_2_ prices, commercial feasibility can be reached independent of the H_2_ price (**Figure 4C**). With that, the AA total market of 25 billion USD can be tackled. BCAAs make a share of roughly 7.3% of the total market with roughly 1.75 billion USD (**Figure 4B**). While this process presents leucine as the first target for commercialization, also valine and isoleucine cell factories can be generated with the presented methods. Dependent on the total bioreactor capacity, this process can supply up to 21% of the total leucine production worldwide using 9 times 500 m^3^ bioreactors (4500 m^3^) Bioreactors of 500 m^3^ are in use in gas fermentation for ethanol production by LanzaTech Inc.^50^.

In conclusion, *M. marburgensis* is a suitable production platform for AA production and, prospectively, other commodity chemicals from CO_2_. Our study shows that carbon-neutral AA production from CO_2_ using methanogens is an alternative to conventional, fermentative AA production. The recently increasing availability of tools for genetic engineering of methanogens and in silico modeling approaches for methanogens facilitates the generation of archaeal cell factories. Breakthroughs in microbial CO_2_ fixation capabilities will also enhance this process. With further developments in the field of renewable energy generation and green H_2_ production, a decrease in prices for AA production from CO_2_ is envisioned. We are highly confident that industrial production of nutrients and chemicals through gas fermentation processes with methanogens is now within our grasp.

## Materials and Methods

### Strain, media, and cultivation conditions

We used *Escherichia coli* NEB5α for cloning and storage purposes, which we purchased from New England Biolabs (New England Biolabs, Ipswich, MA, USA). For the conjugation of *M. marburgensis* with *E. coli*, we utilized the plasmid-mobilizing *E. coli* S17-1 (DSM 9097), acquired from *Deutsche Sammlung für Mikroorganismen und Zellkulturen* (DSMZ, Braunschweig, Germany)^51^. We defined *M. marburgensis* Δ*hpt* as our base strain in this study and referenced it as wild type throughout the manuscript^34^. All generated and purchased strains of *E. coli* and *M. marburgensis* are listed in **Table 2**.

**Table 2.** List of all purchased or genetically modified strains of E. coli or M. marburgensis.

| Strain | Function | Reference |
| --- | --- | --- |
| *Escherichia coli* strains |  |  |
| ***E. coli* NEB5α** | Cloning strain | 52 |
| ***E. coli* S17-1** | Conjugation strain | 51 |
| ***M. marburgensis* strains** |  |  |
| **$\Delta hpt$** | base strain of *M. marburgensis* for marker-less mutagenesis with *hpt* negative selection | 34 |
| **$\Delta hpt$ B4** | mutagenized IPMS for leucine production strain | This study |
| **$\Delta hpt$ C4** | mutagenized IPMS for leucine production strain | This study |
| **$\Delta hpt$ C5** | mutagenized IPMS for leucine production strain | This study |
| **$\Delta hpt$:AHASmt_native** | AHAS overexpression strain | This study |
| **$\Delta hpt$:AHASmt_native E3** | AHAS overexpression strain with mutagenized IPMS for leucine production | This study |
| **$\Delta hpt$:AHASmt_native E4** | AHAS overexpression strain with mutagenized IPMS for leucine production | This study |
| **$\Delta hpt$:AHASmt_native A7** | AHAS overexpression strain with mutagenized IPMS for leucine production | This study |

We cultivated *E. coli* in liquid Luria Bertani broth (LB broth) (Carl Roth GmbH + Co. KG, Karlsruhe, Germany) at 37°C with 180 rpm in a shaker incubator (Labwit, Burwood East (VIC), Australia). We added appropriate concentrations of chloramphenicol (30 µg mL^-1^), ampicillin (100 µg mL^-1^), or trimethoprim (10 µg mL^-1^) for selection to the media when required. For solidified media plates, we also incorporated 1.5% (w/v) of Kobe Agar (Carl Roth) into the LB broth and incubated it at 37°C in a static incubator (Binder GmbH, Tuttlingen, Germany).

We cultivated *M. marburgensis* under the same conditions as described in ^34^. For conjugation experiments and outgrowth thereof, we used the sea salt media (MS media) formulation^35^, and for phenotypic characterizations and bioreactor fermentations, the designated *M. marburgensis* media formulation^53^. Both formulations were reduced by 0.5 mol L^-1^ Na_2_S · 9H_2_O. Closed batch experiments up to 20 mL and 96-deepwell plate experiments contained 4 mL L^-1^ of a 0.025% (w/v) resazurin sodium salt solution as a redox indicator. Where necessary, we added 250 µg mL^-1^ of neomycin for selection of genetically modified *M. marburgensis*. We added 1.5 % Bacto™ agar (Becton, Dickinson and Company, Franklin Lakes, NJ, USA) to solidified media plates and 2.5 g L^-1^ of peptone and 1.25 g L^-1^ of yeast extract to conjugation plates for spot mating.

We performed high-throughput screening in 96-deepwell plates by addition of 2 mL of liquid, sterile reduced MS media to each well, and added individual colonies per well with sterile pipette tips. We sealed the plates with a Breathe-Easy® gas-permeable foil (Merck Group, Darmstadt, Germany) to avoid evaporation. We prepared the plates in an anaerobic chamber (Coy Laboratory Products, Grass Lake, MI, USA) and transferred them into a custom-made stainless-steel anaerobic jar, where we subsequently exchanged the gas atmosphere to H_2_ and CO_2_ (20 Vol.-% CO_2_ in H_2_). We incubated anaerobic jars routinely at 62°C in a static incubator. We usually observed sufficient growth after 36 h of incubation.

We did closed batch phenotypic characterizations with 25 mL of *M. marburgensis* media in 120 mL serum bottles with the trace element composition given in the paragraph for fed batch cultivation and 2 bar overpressure of H_2_/CO_2_ (20 Vol.-% CO_2_ in H_2_) as broadly discussed in^34^. We generated overnight cultures of the respective *M. marburgensis* mutant strains. From those, we inoculated day cultures in biological triplicates to an OD_600_=0.05. We incubated the triplicates at 62°C, shaking at 170 rpm (Eppendorf New Brunswick Innova, Hamburg, Germany). We measured the OD_600_ and amino acid concentrations every 3 h for at least 9 h of incubation and after an additional overnight incubation with a spectral photometer (Hach Lange GmbH, Berlin, Germany) and high-pressure liquid chromatography with mass spectrometry (HPLC-MS) (see amino acid quantification with HPLC-MS).

### Seed inoculum train

We inoculated 25 mL of *M. marburgensis* media from anaerobic glycerol stocks of the respective strain and cultivated overnight at 65°C to reach an OD_600_ of ∼0.3. With this, we inoculated 0.5 L *M. marburgensis* media in 1 L pressure plus bottles (Schott) to an initial OD of 0.01 and cultivated with repetitive regassing to 1 bar overpressure of H_2_/CO_2_ (volume 20% H_2_ in CO_2_) to OD_600_ of ∼1. This was a sufficient cell density to allow for an initial OD_600_ of 0.1 as start density in 1.6 L and 13 L fermentations. For 100 L fermentations, we generated inoculum with a 13 L batch fermentation prior to use.

### Fed batch cultivation

The organism was cultivated in fed-batch mode in three different reactor systems, with varying working volume, such as 1.6 L (DASGIP® 2.2 L bioreactor systems; SR1500ODLS, Eppendorf AG, Hamburg, Germany), 10 L (NLF 19L/13L, Bioengineering AG CH 8636 Wald), and 100 L (P100 F2 150L/100L, Bioengineering AG CH 8636 Wald).

Before inoculation, the reactor was sterilized with the respective working volume of demineralized water containing 3.4 g L^-1^ KH_2_PO_4_ to provide enhanced buffer capacity and phosphorus source. After the sterilization, the liquid was gassed with nitrogen gas to ensure anoxic conditions and we subsequently added 6 mL L^-1^ of the trace element stock solution defined as NSTE (Haslinger, Reischl *et al*. submitted for publication) (containing 90 g L^-1^ Nitrilotriacetic acid, 2 g L^-1^ MgCl_2_ · 6H_2_O, 10 g L^-1^ FeCl_2_ · 4H_2_O, 0.01 g L^-1^ CoCl_2_ · 6H_2_O, 2.4 g L^-1^ NiCl_2_ · 6H_2_O, 0.01 g L^-1^ NaMoO_4_ · 2H_2_O), 2.3 mL L^-1^ of a 30% NH_4_OH solution as a nitrogen source and pH control, and 4 mL L^-1^ of 0.5 mol L^−1^ Na_2_S · 9H_2_O to supply elemental sulfur and regulate the redox potential. Standard fermentation conditions were 60°C, pH 7.0, and vessel volumes per minute (vvm) of 0.25 H_2_/CO_2_ (4:1). Stirring speed depended on the reactor scale and was set to 1500 rpm, 1000 rpm, and 750 rpm for 1.6 L, 10 L, and 100 L, respectively. As high rotation speed is not feasible at large scale, the 150 L reactor was pressurized to 0.5 bar overpressure during fermentation to ensure sufficient gas availability. All other systems were operated at ambient pressure. The gas flow was regulated by the mass flow controllers internal to the system, an external MFC for H_2_ gassing had to be applied for the 1.6 L bioreactors (SIERRA SmartTrak C100L, Sierra Instruments, Monterey, USA).

Inoculation volume varied between 1 and 10% of the working volume, depending on the optical density of the culture. Growth was monitored by regularly measuring the optical density at 578 nm (UV/VIS Spektralphotometer DR 6000, HACH LANGE GmbH, Vienna, Austria) for the small-scale systems or 600 nm (IMPLEN OD600 DiluPhotometerTM, IMPLEN GmbH, Munich, Germany) for the 150 L system. An additional indicator of cell activity was the formation of methane in the gas phase, measured using in-line gas sensors for H_2_, CO_2_, and methane (Sensors by BlueSens gas sensor GmbH, Herten, Germany, as well as AWITE BIOENERGIE GmbH, Langenbach, Germany). At exponential growth, we started feeding an additional shot of trace elements, Na_2_S, and NH_4_OH to avoid liquid limitation. Liquid additions were made via peristaltic pumps (DASGIP MP4/MP8; Biocomponents Peripex W1/W2). To counteract the alkalinity of ammonium hydroxide, phosphoric acid was used for pH control. Redox potential and pH values were monitored by individual probes (Mettler Toledo GmbH, Wien, Austria). Antifoam (Struktol SB2023, Schill und Seilacher, Hamburg, Germany) was added on demand.

We took at each time point liquid samples of 2 mL for amino acid quantification and centrifuged the samples at 16100 g for 15 min at room temperature (5415 R, Eppendorf AG, Hamburg, Germany). The supernatant was transferred to sterile reaction tubes and stored at -20 °C for further analysis.

Fed-batch cultivation was stopped at stationary growth phase, meaning no major increase in either OD_578_ or methane production. This was typically reached after three days of fermentation.

### Continuous fermentation

We performed continuous cultivation fermentations in a 1.6 L working volume. Before we transitioned to the continuous cultivation mode, we enriched the respective *M. marburgensis* strains to suitable cell densities of 1 gCDW L^-1^ with in-situ fed-batch cultivation as discussed above. We defined default process parameters at the start of continuous fermentation as 65°C, pH 7.0, stirring speed of 1500 rpm, a dilution rate of 0.025 h^-1^, and vvm of H_2_/CO_2_ (4:1) of 0.25 at ambient pressure (Haslinger, Reischl *et al.*, submitted). We continuously fed *M. marburgensis* media reduced to 6.8 g L^-1^ KH_2_PO_4_, 2.1 g L^-1^ NH_4_Cl as nitrogen source, and trace elements. The concentration and composition of the trace element stock solution depended on the experimental set-up and either corresponded to those mentioned above or were reduced to 1 mL L^-1^ with different metal concentrations defined as OriTE (40 g L^-1^ MgCl_2_ · 6H_2_O, 10 g L^-1^ FeCl_2_ · 4H_2_O, 0.2 g L^-1^ CoCl_2_ · 6H_2_O, 1.2 g L^-1^ NiCl_2_ · 6H_2_O, 0.2 g L^-1^ NaMoO_4_ ·2H_2_O) or NSTE (see above). The media for continuous feeding was prepared in sterile Nalgene containers (20 or 50 L Nalgene™ Polypropylene, Carboy) by mixing concentrated and autoclaved salt solution with filter sterilized RO water. To mitigate foaming, we added 100 µL L^-1^ antifoam. The mixture was then flushed for 30 min with nitrogen gas to ensure anoxic conditions before feeding them to the reactor. 0.5 mol L^−1^ Na_2_S · 9H_2_O solution as sulfur source and to control the redox potential were adjusted to the dilution rate. To keep the volume steady and prevent liquid overflow we adjusted the height of the harvesting pipe and continuously pumped fermentation broth into a second Nalgene container. Liquid sampling and monitoring optical density and methane formation was done as described above. Steady state was considered after 3 volume exchanges and steady growth behavior.

### Data interpretation

The amino acid concentration (g L^-1^) measurements built the base for further calculations, such as the volumetric productivity (g L^-1^ h^-1^) (**Equation 1**) or specific productivity (g gCDW^-1^ h^-1^) (**Equation 2**), while gCDW is depicted as g^-1^ throughout the results part. It requires at least 2 timepoint measurements and is defined as in **equation 3**.

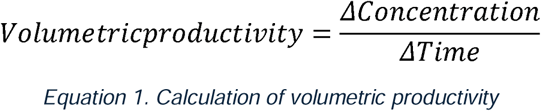

The specific productivity is then defined as in equation 2:

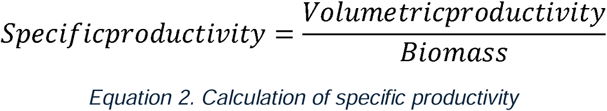

We chose to approximate the biomass (gCDW) as the median point between the starting biomass and the final biomass. This underestimates the average biomass and could slightly overestimate the specific productivity in the exponential phase. So, the biomass is defined as in **equation 3** with *a* factor to convert OD_600_ in grams of cell dry weight equal to 0.371^54^.

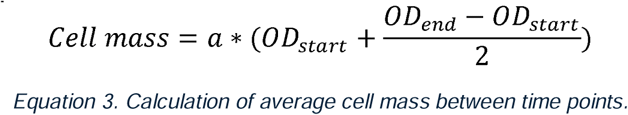

### Molecular cloning techniques

All primers were purchased from Eurofins genomics (Eurofins Genomics Europe Shared Services GmbH, Ebersberg, Germany) and are listed in **Supplemental Table 2**. The synthesized fragment of the codon-optimized acetolactate synthase (AHAS0) was synthesized by Twist Biosciences (South San Francisco, CA, USA) and delivered in high copy number ampicillin resistance standard vector (**Supplemental Table 3**).

In the first step, we utilized the pArk00005 as a base vector and integrated *Xba*I as additional unique restriction enzyme recognition site between *Avr*II and *Kas*I^34^. With that, we created two additional modules between *Avr*II and *Kas*I resulting in pArk00007. Afterwards, we exchanged the non-sense DNA between *Kas*I and *Xba*I with the Twist Bioscience synthesized fragments of AHAS0 resulting in pArk00008. We purchased all necessary restriction enzymes and the sticky-end ligation kit from New England Biolabs (Ipswich (MA), USA). We used chemically competent NEB5α for standard cloning approaches and storage and chemically transformed *E. coli* S17-1 with a confirmed plasmid with Sanger Sequencing for consecutive conjugation with *M. marburgensis*.

### Transformation of *M. marburgensis*

We carried out the transformation of *M. marburgensis* via conjugal DNA transfer from *E. coli* S17-1 to *M. marburgensis* on a spotmating-based setting as previously described in Klein et al. 2025. We cultivated *M. marburgensis* and plasmid-carrying *E. coli* to the late exponential growth phase, centrifuged ∼2·10^8^ cells of *M. marburgensis* and ∼2·10^9^ *E. coli* cells anaerobically and mixed the pellets in sterile anaerobic MS medium inside the anaerobic chamber. We spotted the cell suspension on an LB/MS solidified media plate and let the spot be absorbed. Afterwards, we incubated the plate with 1 bar overpressure of H_2_/CO_2_ (20 Vol.-% CO_2_ in H_2_) in a stain-less steel jar for 18 h at 37°C. Afterwards, we washed off the spots with 1 mL of sterile MS media and recovered the transformants in 5 mL of MS media for 4 h at 65°C. 1 mL of recovered *M. marburgensis* was subsequently transferred to 250 µg mL^-1^ neomycin-containing MS medium for liquid selection of positive transformants. We isolated monoclonal transformants from selective purity plating and confirmed the purity of the strain with PCR-based analysis.

### PCR-based analysis of positive transformants

After two selective transfers or purity plating, positive transformants were analyzed towards double homologous recombined target specific integration of AHAS0. Therefore, we tested for integration with four specific primer combinations (Seq_Ark_12+06; Seq_Ark_02+05; Seq_Ark_05+06; Seq_Ark_23+05) to elaborate on single homologous or double homologous recombination and wild-type background. If wild-type background was still observed, we performed a second purity plating to isolate monoclonal strains with double homologous recombination of AHAS0. The PCR follows a colony PCR-like protocol following the Phire plant mastermix manufactureŕs manual (Thermo Scientific, Waltham (MA), USA)^34^.

### EMS treatment + 4-azaleucine selection

For random mutagenesis of *M. marburgensis*, we took 5·10^8^ late exponential cells, concentrated with sterile MS media to 2.5·10^8^ cells mL^-1^ via centrifugation at 5000 g for 4 min at room temperature in the anaerobic chamber. We added EMS solution to a final concentration of 0.1 mol L^-1^ and incubated the sample for 7.5 min at 37°C in a static incubator. The concentration and incubation time were experimentally defined through a screening experiment to achieve a survival rate of approximately 50% (**Supplemental Table 4**, **Supplemental Figure 8**). Afterwards, we added 6% w/v of thiosulfate solution to the mutagenized cells and mixed the cell suspension thoroughly. Furthermore, the cell suspension was washed twice with sterile MS media in the anaerobic chamber with centrifugation conditions as described above and finally resuspended in 1 mL of sterile MS media. From this mutagenized *M. marburgensis* we generated anoxic glycerol stocks with a final glycerol content of 25% (w/v).

We selected leucine overproduction mutants using 4-azaleucine, as it has been reported to bind allosteric leucine binding sites with higher affinity and results in inhibition of leucine biosynthesis. Mutations in the allosteric binding sites of leucine prohibit the binding of 4-azaleucine/leucine and result in allosteric release and overproduction of leucine ultimately. Therefore, we prepared solidified MS agar plates containing 1 mg mL^-1^ of 4-azaleucine and plated 10^7^ cells of the mutagenized *M. marburgensis* mutant library. We incubated the plates at 62°C in a stainless-steel jar with 1 bar overpressure of H_2_/CO_2_ (20 Vol.-% CO_2_ in H_2_). Colony formation was usually found after 3 days of incubation. We transferred colonies to 96-deepwell plates as described above and analyzed leucine concentrations after substantial growth occurred (OD_600_>0.1).

### Sequencing analysis of leucine production mutants

In the first step, we PCR amplified the IPMS and the AHAS genes of leucine overproduction mutants and sequenced the respective genes with Sanger sequencing (Seq_Ark_21+22; Seq_Ark_23+24; Seq_Ark_25+26) at Eurofins Genomics (Eurofins Genomics Europe Shared Services GmbH, Ebersberg, Germany). The sequencing results were analyzed with Snapgene®. Ultimately, we prepared multi-sequence alignments of the truncated *imps* gene sequences in comparison with wild-type *M. marburgensis ipms* sequences and allosteric inhibition-released IPMS-encoding gene versions from *E. coli* and *C. glutamicum*^43^. The multisequence alignment was generated with Clustal^55^.

### In silico analysis of IPMS structure and identification of leucine binding sites

We compared the structure of the IPMS enzyme from *M. marburgensis* with the corresponding enzymes from *E. coli* K12 and *C. glutamicum in-silico*. Therefore, we gathered the enzyme structures predicted by alpha-fold 2 from Uniprot^56^. Afterwards, we performed an alignment analysis in PyMol^57^. To identify putative leucine binding sites in the IPMS enzymes, we transformed leucine.sbf to leucine.pdb in PyMol and uploaded it into Autodock as ligand. Additionally, the three IPMS enzyme structure predictions were uploaded to Autodock. The water bonds were removed, Kollman charges were added, and polar H_2_ connections were added. Afterwards, we defined the relevant area of the enzyme for the leucine binding sites. The resulting config file (**Supplemental Figure 9**), including the IPMS enzyme and leucine as ligand, was read into Autodock Vina for binding site prediction based on kCal Mol^-1^ binding affinities^58^. The 10 highest binding affinities were given as output. The output of each enzyme from *M. marburgensis*, *E. coli*, and *C. glutamicum* IPMS for the most likely leucine binding site was added to PyMOL and overlayed with the respective enzyme structure. Ultimately, all predicted leucine binding sites and enzyme structures of the three species were aligned and exported as figure.

### Crude extract preparation for enzyme assays

We regularly generated crude extract cell lysate of *M. marburgensis* for enzyme assays. Therefore, we stepwise centrifuged 10-14 mL of late exponential *M. marburgensis* from closed batch cultivation with OD_600_=0.35 for 4 min, 4500 g, at room temperature in the anaerobic chamber. We resuspended the cell pellet in 0.5 mL of water. In the next step, we bead beaded the cells with 0.1 cm zirconium beads two times 20 s with 20 s of break in between (Fastprep-24, MP Biomedicals, Irvine, CA, USA) To separate zirconium beads and cell debris, we centrifuged 5 min, 13000 g, at room temperature and anaerobically transferred the supernatant to a sterile reaction tube.

### Acetolactate synthase (AHAS) enzyme assay and analysis of leucine and valine allosteric inhibition

All initial steps involving the enzymatic reaction of AHAS were carried out in the anaerobic chamber since *M. thermautotrophicus* AHAS turned out to be a strictly anaerobic enzyme as it has been demonstrated for *M. maripaludis* before^41^. The assay buffer contained 20 mmol L^-1^ sodium pyruvate, 0.5 mmol L^-1^ TPP, 10 μmol L^-1^ FAD, and 10 mmol L^-1^ MgCl_2_ in 50 mmol L^-1^ Tris buffer (50 mmol L^-1^ Tris-HCl, 150 mmol L^-1^ NaCl, pH=7.0) and we anaerobized it with N_2_/CO_2_ (20 Vol.-% CO_2_ in N_2_) for 30 min on ice. We mixed 20 µL of assay buffer with 2 µL of *M. marburgensis* crude extract in 0.5 mL PCR tubes. The tubes were anaerobically incubated for 1 h at 60°C. Thereafter, we added 2 µL of 4 mol L^-1^ sulfuric acid, mixed well, and incubated at 60°C to convert acetolactate to the more stable acetoin by spontaneous decarboxylation. To develop the characteristic red color for the colorimetric assay, we mixed creatine (0.5% (w/v)) and naphthol (5% naphthol (w/v) in 2.5 mol L^-1^ NaOH) in a 1:1 ratio and added 36 µL of the mixture to the sample. After further incubation at 60°C for 15 min, we centrifuged the samples for 5 min, 13000 g, at room temperature and measured the supernatant absorbance at 525 nm using a 384-well plate with a Tecan Spark plate reader (Tecan Group Ltd, Maennedorf, Switzerland).

To define allosteric inhibition of AHAS by valine or leucine, we added 0 to 25 mmol L^-1^ of valine or leucine, respectively, to the assay mix and compared to outcome to controls without leucine or valine addition.

### Amino acid quantification with HPLC-MS

All 20 proteinogenic amino acids were quantified directly from the growth medium. In brief, after the removal of cells, the supernatant was diluted in acetonitrile (1:3) and was warmed up for 5 min to 40°C to facilitate protein precipitation. After centrifugation (14 000 g; 5 min), the supernatant was mixed with the IS (4:1; IS was 10 µmol L^-1^ of Metabolomics Amino Acid Mix purchased from Cambridge Isotope Laboratories, Inc.). For the chromatographic separation the LC system (Agilent 1260 Infinity II Prime LC) was coupled to a MS detector (Agilent 6475 Triple Quadrupole) and separation was achieved with a Poroshell 120 HILIC-Z analytical column (2.7 µm, 2.1·100 mm) together with gradient elution from 90/10 to 65/35 25 mmol L^-1^ aq. NH_4_COO/MeCN (pH=3). Analytes where ionized using ESI in positive mode and analyzed with dynamic MRM. A 9-point external calibration curve with concentrations from 76 nmol L^-1^ to 125 µmol L^-1^ was used for quantification.

### Techno-economic model and sensitivity analysis

A techno-economic model including a sensitivity analysis was created for process evaluation and to better predict the variables in our process. The model was created based on gas fermentation using the main synthesis pathways. Input materials such as H_2_, CO_2_, NH_3_, and the micro and macro elements used were taken into account.

The total was balanced with the outgoing elements that flow into the AAs, methane, and biomass (**Supplemental Material 3, Supplemental Figures10+11**). The elemental composition of the respective substances is decisive here.

The model is flexible in this respect, as we can modify it based on the organism’s production- and productivity-specific data. The costs for raw materials and other OPEX, as well as CAPEX, can also be entered as input parameters. The different CAPEX and OPEX costs can be further divided into seven different process steps (H_2_ production, resources, fermentation, gas utilization, DSP, packaging, utility) so that the costs can be allocated proportionally to the process areas. Since three different products are created from the process, the specific production costs can be represented by different allocations.

Based on this structure of the TEA, it is possible to simulate different scenarios by varying process and production parameters and to evaluate them based on the influence of the parameters on production costs. The resulting sensitivity analysis provides guidance for prioritizing the further development of the process. The price of H_2_ and the MER/LVPR ratio were identified as the main influencing factors. Other important influencing factors are CO_2_ costs, LVPR, and MER. The sensitivity analysis process was carried out by assuming different minimum and maximum values for the individual areas. These were then subdivided into values in this range of “good,” “bad,” and average. For the sensitivity analysis, all scenarios with different values were examined. One process parameter was varied, and the remaining values were set to average. This allowed an iterative ranking to be developed with the process parameters that were analyzed as having the highest impact on process costs.

An evaluation of current and projected H_2_ prices was carried out based on data provided by the International Energy Agency (IEA), assuming cost scenarios of 2 € kg^⁻¹^ and 4 € kg^⁻¹^ for grey H ^59^. These values served as a foundation for the subsequent techno-economic analysis of the proposed bioprocess. The process model was established for a reactor capacity of 1000 m³ per bioreactor unit, representing an industrially relevant scale. The biomass selling price was estimated with 0.50 € kg^⁻¹^.

## Supplementary materials

The raw data will be uploaded to the PHAIDRA repository system of the University of Vienna. DOI will be displayed here.

## Author contributions

C.F. and F.S. designed the laboratory experiments; F.S., A.R., E.H., S.D. and S.G. performed process development and operated the bioreactors; C.F., A.R. and M.K performed genetic engineering of *M. marburgensis*; S.D. and F.U. performed standard cloning methods; F.U. performed enzyme assays and phenotypic characterizations; C.F.,, F.S, R.F., J.S., G.B. and S K-M. R. R.. supervised the project; G.B. and S.K-M.R.R. acquired funding; F.S.,J.S., S.S. and G.B. designed the 150 L bioreactor *Seestadt* facility container lab; G.B. and B.A. designed the TEM with input from C.F., J.S. and S.K-M.R.R., B.A. and G.B. performed the sensitivity analysis; T.S-P. analyzed the AA samples; C.F., E.H., G.B. and S.K-M.R.R. wrote the manuscript, while all authors edited the final manuscript and approved its publication.

## Acknowledgements

The COMET center: acib: Next Generation Bioproduction is funded by BMIMI, BMWET, SFG, Standortagentur Tirol, Government of Lower Austria und Vienna Business Agency in the framework of COMET - Competence Centers for Excellent Technologies. The COMET-Funding Program is managed by the Austrian Research Promotion Agency FFG. Open access funding was provided by the University of Vienna.

