## Supplementary material for "Pilot-scale production of leucine from CO_2_"

**Running Head: Leucine Production *M. marburgensis***

### Supplemental materials and methods

**Supplemental Material 1: Primers, Plasmids, and synthesized DNA fragments**

Supplemental table 1. Plasmid list.

| Plasmid constructs | Function | Reference |
| --- | --- | --- |
| pArk00005 | Basis vector for genome integration in *M. marburgensis* | ^1^ |
| pArk00007 | Basis vector with modular integration site for genome integration at *hpt* | This study |
| pArk00008 | Native acetolactate synthase containing vector for integration in *M. marburgensis.* | This study |
| TWIST_AHASmt_native | TWIST Bioscience vector with acetolactate synthase from *M. thermautotrophicus* codon-optimized for *M. marburgensis*. | This study |

Supplemental table 2. Primer list.

| Name | Purpose | Sequence (5' --> 3') | Reference |
| --- | --- | --- | --- |
| Seq_Ark_05 | specific for gDNA *M.m.* *hpt* region downstream flank | CCACCGATCCCAGGTAAGATCG | ^1^ |
| Seq_Ark_06 | specific for gDNA *M.m.* *hpt* region upstream flank | CCTCAACCTTTTCAAGGAGGCTC | ^1^ |
| Seq_Ark_12 | Inside *hpt* gene | GGGATGTCCTCAACGGTGACC | ^1^ |
| Seq_Ark_01 | specific for pArk Vector system | CAGGAAACAGCTATGACCACTAG | ^1^ |
| Seq_Ark_02 | specific for pArk Vector system | CAACTTGCCCACTGCTAGC | ^1^ |
| Seq_Ark_22 | FW Sequencing primer for AHAS from *M. thermautotrophicus* (*ilvB*) | CCGTGACAACAACACTCCTTGG | This study |
| Seq_Ark_23 | FW Sequencing primer for AHAS from *M. thermautotrophicus* (*ilvB*) | GTCCCATACACACCTGGGCG | This study |
| Seq_Ark_21 | RV Sequencing primer for AHAS from *M. thermautotrophicus* (*ilvB*) | CCGTGCATGCCGAGCATAC | This study |
| Seq_Ark_24 | RV Sequencing primer for AHAS from M. thermautotrophicus (*ilvN*) | CCATGCAGAGCTCCCTCTTAAC | This study |
| Seq_Ark_25 | FW Sequencing primer for AHAS from *M. marburgensis* (*ilvB*) | CCGGGCGAAACATCTGAAGC | This study |
| Seq_Ark_26 | Sequencing primer for AHAS from *M. marburgensis* (*ilvB*) (50 bp upstream) | CCTCCTCATCTCTAATTAACAGGG | This study |
| Seq_Ark_27 | Sequencing primer for IPMS from *M. marburgensis* | GCCCGAAGATCTGCGTCG | This study |

Supplemental table 3. List of synthesized DNA fragments.

| Name | Insert sequence | Reference |
| --- | --- | --- |
| AHAS 0 codon optimized | TCTAGAAAATGTCCTGAAATAAAAAAAATTAGGTGCATTACACTTAAAAAGTTTGTGCAATGCACCTTCGTTAATCAGTATCGGCTCACATTGTCCTGCTTCCCCTGGACATTGCTGTTGGCCCTGTCCTTGCGAGTTCCTTTATTCCAAAGTTCCTCAGAAGCTCCAGGAAGGCGTCTATCTTTTCTGAGTCCCCTGTAACCTCAACTGTCAGGGCATCGGGTGACACATCCACTATCCTCCCCCTGAATATGTTGGTGTACTGTATTATCTCTGACCTTTCTGACTCTGATGGGGCGTGGACCTTGACCATGCAGAGCTCCCTCTTAACTGTTGCTGCGGGTTCAAGGTCCCTGACCTTTATGACGTCTATGAGCTTGTTCAGCTGCTTTGTTATCTGCTCAAGCACCCTGTCGTCGCCCCTGGCGATTATTGTCATCCTGGCAATGCCTGGGGTTTCTGATTCCCCGACTGTTATGTTTTCAATGTTGAATCCCCTTCTTGTGAAGAGTCCTGCAACCCTCTGGAGCACTCCTGGTTTGTGCTCCACGAGGGCGCTTATGATATGGGTATCGGGTTCCATCTCAATCACCATCCGCCTTCCCTGGGGAGTATTTAATTTCCTGGGGGTCCTCCCTTTCAACCCTGTACTCCCCCACTATCTCGGTGAGACCACAGCCGGGGGGGACCATGGGGAGTATCTCATCAGGATCTATCACTATATCAAGGAGGGCAGGTTCACCTGACCTTATTGCCCTTGAAAGGGCTTCTGAGGTTTCACCGGGTTCCTCTATCCTCTCTGCCTCCACTCCAAATGATTCTGCCAGCTTCACAAAGTCGGGAACTTCGCCCAGGTGTGTATGGGACATTCTCTCATCATAGAAGAGCCTCTGCCACTGTGCCACCATTCCAAGGTGCCTGTTGTCCATGATACATATCACCACGGGGATGTCGTATTCCCTTATGGTTGCAAGGTCCTGGCAGACCATGAGGAATCCGCCGTCACCGCACACTGCAACAACGTCTGAATCAGGCAGTGCCACCTTGGCACCTATGGCGGCTGGAAAACCGAAGCCCATTGTTCCAAGGCCTCCTGATGATATGAACTTTCTGGGGGCCCTGGATGTGTAGAAGTGGGCCATCCACATCTGGTTCTGTCCCACATCTGTTGTAACGACTGTCTCGTCATCAAGGACCTGGCTTATCTCCTTTATAACCTGCTGGGGCTTCAGGGGCACCTCATCATAGCTCATCCTTGGCATGCAATCGGCCCTGAATTTCTGGACGCTTTCAAGCCACTGGCTGTCCCTCTTTTCATATTTTTTGAGTTTTGCTATGAGTTCCCTGAGGACGTTTCTTGCATCTCCAACGATGGGGACATCAACCCCAACGTTCTTACCTATCTCTGCGGGGTCGATGTCGACGTGTATTATCCTGGCGTTGGGGGCGAATTCTGCAACGTTCCCTGTTGTCCTGTCTGAGAATCTGCATCCAACGGCTATGAGGCAGTCGCATTCGTCCACTGTCAGGTTTGCCACCTTCCTGCCGTGCATGCCGAGCATACCCATGGCTGAAGGGTGGTCCTCAGGAAAGGAACCCTTACCAAGGAGTGTTGTTGTCACGGGGGCCTTTATGAGATCTGAGAGTTCCTTTATCTCCCTGGATGCCCCTGATATTATAACTCCTCCACCTGCAAGTATGACGGGTTTTTCTGACCTCCTTATGAGTTCTGCGGCCCTCTTTATCTGGAGGGGGTGGCCCTTAACATTGGGCCTGTACCCTGGGAGCTCCAGGTCATCAACCTCCTCCATGATCTCCTGTTCCTGTATATCCTTGGGGAGGTCTATAACAACGGGTCCTGGCCTTCCTGTCTTTGCTATGTGGAAGCTTGCCCTGACAATTGCAGGTATCTCGCTGGCGTCTGATGGCTGGAATGAGTGCTTGGTGATGGGCATGGTTATCCCTATCATGTCCACCTCCTGGAATGCATCATTTCCAATGAGGTGTGTTGGGACCTGACCTGCAATGGCCACGATGGGGGCTGAGTCCATGTAGGCTGTTGCAATGCCTGTAACAAGGTTTGTTGCCCCGGGACCGGAGGTTGCTATGCAGACCCCCACCCTTCCTGAGGCCCTTGCATATCCGTCTGCTGCGTGTGCTGCGCACTGTTCATGTCTAACGAGGATGTGTTTAAGTTCTGAATCATAGAGCATATCATAGAGTGGCAGGAGCTGTCCACCGGGGTATCCGAAAACGGTGTCTGCTCCCTGATCCAGAAGTGATCTGATTATTGCCTGGCCACCTTTCATTGGAAACACAATTAACCACCTCATTTTATGTGATATTATCTATTCATATAATCCTATATAAATATATCGCTAATTTTAAGGTTTTTCTGAGCCATCGGTTGGTTCATGGGGGCGCC | This study |

**Supplemental Material 2: Definition of EMS treatment survival rate of *M. marburgensis***

First, we defined the plating efficiency of *M. marburgensis* on solidified agar plates (**Supplemental Table 4**). Based on that, we defined the suitable concentration and incubation time of 1 mol L^-1^ EMS and 7.5 min for prospective random mutagenesis experiments by generation of a kill curve (**Supplemental Figure 8**). After the random mutagenesis, we plate-selected for leucine overproducers by the addition of the competitive toxic leucine analog 4-azaleucin and tested 25 mutants for overproduction of leucine compared to wild-type *M. marburgensis*.

**Supplemental Material 3: Generation of carbon balances**

Carbon balances were calculated based on the C-mol carbon that entered the bioreactor as CO_2_. This C-mol carbon was balanced against carbon in the off-gas as CO_2_ or biomethane (CH_4_) or as carbonated substances in the fermentation broth, such as amino acids, biomass, other organic acids, or unknown substances. The C-mol carbon of biomass was deduced from previous studies^2^

**Supplemental results**

Amino acid percentage / %

| Mutant identifier | Alanine | Glutamic acid | Glycine | Isoleucin | Leucine | Valine |
| --- | --- | --- | --- | --- | --- | --- |
| A1 | 59.3205% | 9.55005% | 8.26446% | 10.7438% | 5.92287% | 6.19835% |
| A2 | 59.8115% | 8.61663% | 9.72736% | 10.6361% | 5.35173% | 5.85661% |
| A3 | 59.8074% | 8.77318% | 7.59629% | 10.7703% | 7.81027% | 5.24251% |
| B1 | 59.2276% | 8.74915% | 9.22761% | 11.1757% | 5.7758% | 5.84416% |
| B2 | 59.5617% | 9.06747% | 8.8526% | 10.6145% | 6.44607% | 5.45767% |
| B3 | 59.6838% | 9.78261% | 7.90514% | 10.7708% | 5.92885% | 5.92885% |
| C1 | 59.4655% | 9.57684% | 6.4588% | 10.5791% | 9.02004% | 4.89978% |
| C2 | 59.5849% | 9.03541% | 7.08181% | 10.6227% | 8.91331% | 4.7619% |
| C3 | 60.1093% | 9.01639% | 7.54098% | 10.765% | 8.03279% | 4.53552% |
| D1 | 59.5769% | 10.8462% | 7.96154% | 10.3462% | 6.76923% | 4.5% |
| D2 | 59.129% | 9.46399% | 6.70017% | 10.9715% | 8.96147% | 4.77387% |
| D3 | 38.1549% | 16.6856% | 7.91572% | 10.0797% | 17.6538% | 9.51025% |
| A4 | 59.2183% | 8.77581% | 6.71091% | 9.88201% | 11.5044% | 3.90855% |
| A5 | 51.7544% | 7.86341% | 6.20301% | 8.45865% | 22.5564% | 3.16416% |
| A6 | 44.8052% | 23.0519% | 11.6883% | 7.79221% | 4.54545% | 8.11688% |
| B4 | 46.2838% | 7.54505% | 4.95495% | 8.44595% | 30.1802% | 2.59009% |
| B5 | 59.2771% | 7.46988% | 11.3253% | 10.3614% | 5.54217% | 6.0241% |
| B6 | 55.4565% | 10.2833% | 9.91605% | 11.49% | 6.55824% | 6.29591% |
| C4 | 46.4015% | 7.38636% | 6.25% | 7.76515% | 29.5455% | 2.65152% |
| C5 | 45.4429% | 7.95892% | 5.5199% | 7.31707% | 31.3222% | 2.43902% |
| C6 | 32.7482% | 4.53995% | 3.87409% | 14.4673% | 17.8571% | 26.5133% |
| D4 | 35.0921% | 14.0227% | 4.85127% | 6.9051% | 33.9943% | 5.13456% |
| D5 | 32.0086% | 14.0172% | 4.7261% | 7.78733% | 35.0698% | 6.39098% |
| D6 | 34.4037% | 17.737% | 5.27523% | 10.1682% | 20.8716% | 11.5443% |

Supplemental Figure 1. Amino acid percentage of 96 deep well high throughput screening. Red boxes mark the leucine percentage of mutants that were pursued in follow up experiments.

Amino acid concentration / mg L-1

|  | A1 | A2 | A3 | B4 | C4 | C5 |
| --- | --- | --- | --- | --- | --- | --- |
| Alanine | 8.51759 | 10.1093 | 9.55882 | 6.07101 | 6.56349 | 6.93152 |
| Glutamic acid | 1.37125 | 1.45637 | 1.40219 | 0.989677 | 1.0448 | 1.214 |
| Glycine | 1.18666 | 1.64411 | 1.21409 | 0.649937 | 0.884062 | 0.841964 |
| Isoleucine | 1.54266 | 1.79771 | 1.72139 | 1.10785 | 1.09838 | 1.11609 |
| Leucine | 0.85044 | 0.904543 | 1.24829 | 3.95871 | 4.1792 | 4.77766 |
| Valine | 0.889996 | 0.989877 | 0.837893 | 0.33974 | 0.375057 | 0.372031 |

Supplemental Figure 2. Concentration of selected AA from 96 deep-well results from high-throughput screening with non-EMS treated wild-type and mutagenized cultures selected with 4-azaleucine as a toxic amino acid analog.

*

*

*

Specific AA productivity / mg g-1 h-1

|  | WT | A5 | D4 | D5 |
| --- | --- | --- | --- | --- |
| Alanine | 5.80697 | 6.03984 | 7.30732 | 6.28368 |
| Glutamic acid | 1.55105 | 2.40093 | 3.41265 | 2.43844 |
| Glycine | 0.567456 | 0.637747 | 0.829063 | 0.787805 |
| Isoleucine | 1.00251 | 0.825319 | 0.925465 | 1.01289 |
| Leucine | 0.321559 | 2.30714 | 2.94992 | 2.47596 |
| Valine | 0.945761 | 0.694018 | 0.829063 | 0.656504 |
| Total | 11.1032 | 13.8616 | 17.3718 | 14.7432 |

Supplemental Figure 3. Specific productivity of AA of three respective mutants (n=2) in comparison to wildtype. Error bars mark the standard deviation (n=2). An asterisk mark indicates statistical significance inferred from Student´s t-test with p<0.01.

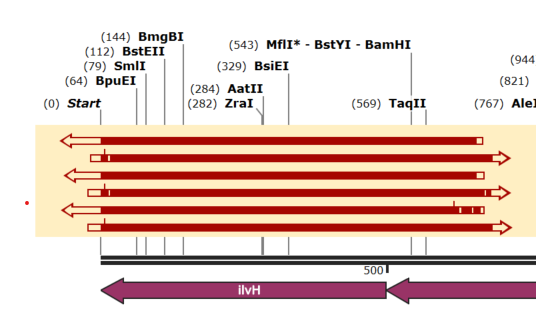

Supplemental Figure 4. Sanger sequencing results of the native regulatory subunit of AHAS, *ilvN* (*ilvH*), in *M. marburgensis* mutants after EMS treatment and 4-Aza-Leucine selection. No mutations were identified. Light red arrows indicate one Sanger sequencing result. All three selected mutants, A7, E3, and E4 were sequenced in both directions.

Amino acid percentage / %

|  | Alanine | Glutamic acid | Glycine | Isoleucin | Leucine | Valine |
| --- | --- | --- | --- | --- | --- | --- |
| A1 | 18.183% | 7.80244% | 2.34423% | 13.497% | 13.3609% | 44.8124% |
| A2 | 18.473% | 7.73519% | 2.18572% | 13.689% | 13.1503% | 44.7668% |
| A3 | 19.834% | 8.42891% | 2.62836% | 13.4244% | 12.647% | 43.0372% |
| A4 | 20.6943% | 8.36558% | 2.51608% | 13.2246% | 12.5898% | 42.6096% |
| A6 | 25.4522% | 4.49664% | 2.66554% | 13.6573% | 16.7946% | 36.9337% |
| A7 | 14.4199% | 3.73148% | 1.50733% | 6.07424% | 58.5837% | 15.6834% |
| A8 | 18.268% | 7.83684% | 1.9791% | 11.1023% | 30.7039% | 30.1098% |
| A9 | 19.6067% | 11.4844% | 2.0824% | 9.60942% | 23.3018% | 33.9152% |
| B1 | 20.4562% | 4.90063% | 2.45548% | 14.3613% | 16.8171% | 41.0093% |
| B2 | 18.7046% | 7.07229% | 2.25467% | 14.018% | 15.9089% | 42.0415% |
| B3 | 25.7441% | 6.13553% | 2.65323% | 14.9794% | 16.3275% | 34.1603% |
| B4 | 17.4741% | 4.05063% | 2.28292% | 12.9773% | 18.2845% | 44.9305% |
| B6 | 17.321% | 5.00605% | 1.69089% | 9.37514% | 31.2935% | 35.3134% |
| B7 | 16.4672% | 4.76275% | 1.75503% | 13.956% | 25.6701% | 37.3889% |
| B8 | 22.3667% | 6.76736% | 2.27424% | 8.82584% | 41.049% | 18.7169% |
| B9 | 20.3512% | 6.67978% | 2.43698% | 7.94852% | 42.3852% | 20.1983% |
| D1 | 20.7934% | 6.80138% | 2.41728% | 14.3001% | 13.5376% | 42.1502% |
| D2 | 21.1874% | 9.79742% | 2.46432% | 13.8815% | 11.6084% | 41.061% |
| D3 | 9.89875% | 8.15159% | 1.37407% | 6.43924% | 36.5043% | 37.632% |
| D4 | 18.9774% | 6.24773% | 2.21384% | 8.93934% | 39.6868% | 23.9349% |
| D6 | 20.5876% | 5.71431% | 2.72492% | 8.21657% | 42.9276% | 19.829% |
| D7 | 14.2029% | 5.12349% | 1.75323% | 8.7062% | 34.914% | 35.3002% |
| D8 | 19.9252% | 3.80344% | 2.11429% | 14.3528% | 19.1147% | 40.6896% |
| D9 | 20.0327% | 5.10213% | 1.9694% | 9.73612% | 38.1469% | 25.0127% |
| E1 | 20.2256% | 3.21595% | 1.75578% | 14.4082% | 18.6119% | 41.7826% |
| E2 | 21.3266% | 7.8289% | 2.26504% | 14.6614% | 13.8434% | 40.0747% |
| E3 | 18.4025% | 4.25636% | 1.9371% | 6.0255% | 56.2339% | 13.1447% |
| E4 | 13.9263% | 2.5712% | 1.38047% | 6.60494% | 50.2401% | 25.2769% |
| E6 | 19.3713% | 4.2245% | 1.76533% | 13.4683% | 19.0596% | 42.111% |
| E7 | 22.0472% | 8.41321% | 2.25507% | 9.97614% | 34.8852% | 22.4231% |
| E8 | 19.1815% | 4.69842% | 2.24431% | 13.0565% | 20.149% | 40.6703% |
| E9 | 23.5442% | 10.4345% | 2.52373% | 8.13574% | 35.3755% | 19.9863% |
| G1 | 19.992% | 6.77188% | 2.17837% | 9.52827% | 35.7349% | 25.7946% |
| G2 | 22.7088% | 9.15367% | 2.50808% | 14.1284% | 12.2581% | 39.2429% |
| G3 | 18.5491% | 7.59163% | 2.21092% | 8.7535% | 43.4045% | 19.4903% |
| G4 | 19.8231% | 8.17761% | 2.19015% | 9.59289% | 39.6706% | 20.5456% |
| H1 | 13.3622% | 3.29153% | 1.0892% | 15.7737% | 15.1047% | 51.3787% |
| H2 | 19.1975% | 8.07275% | 2.28776% | 8.98856% | 40.5647% | 20.8887% |
| H3 | 19.8183% | 6.65555% | 2.19893% | 8.71099% | 41.9602% | 20.6561% |
| H4 | 20.3739% | 7.83389% | 2.62758% | 13.6654% | 18.1611% | 37.3381% |

Supplemental Figure 5. AA percentage of 96 deep-well high throughput screening with AHAS0. The first three colonies derived from non-EMS treated AHAS0. The rest is from EMS treated AHAS0. Red boxes mark the leucine percentage of mutants that were pursued in follow up experiments.

Amino acid concentration / mg L-1

|  | A1 | A2 | A3 | A7 | E3 | E4 |
| --- | --- | --- | --- | --- | --- | --- |
| Alanine | 2.45137 | 2.69326 | 2.26833 | 3.2567 | 3.85584 | 3.4272 |
| Glutamic acid | 1.0519 | 1.12775 | 0.963979 | 0.842747 | 0.891827 | 0.632759 |
| Glycine | 0.31604 | 0.318666 | 0.300595 | 0.340428 | 0.405877 | 0.339726 |
| Isoleucine | 1.81961 | 1.99579 | 1.53529 | 1.37185 | 1.26251 | 1.62544 |
| Leucine | 1.80127 | 1.91724 | 1.44639 | 13.231 | 11.7826 | 12.3638 |
| Valine | 6.04144 | 6.52676 | 4.92198 | 3.54206 | 2.75418 | 6.22053 |

Supplemental Figure 6. Concentration of selected amino acids from 96 deep-well results from high-throughput screening with non-EMS treated AHAS0 mutant and mutagenized cultures of AHAS0 selected with 4-azaleucine as a toxic amino acid analog.

Specific AA productivity mg g-1 h-1

|  | A7avg | E3avg | E4avg |
| --- | --- | --- | --- |
| Alanine | 3.72226 | 3.749 | 4.66191 |
| Glutamic acid | 1.38459 | 1.8934 | 1.9159 |
| Glycine | 0.187135 | 0.165848 | 4.90137 |
| Isoleucine | 4.14697 | 0.970817 | 5.26604 |
| Leucine | 20.1598 | 25.6745 | 19.1727 |
| Valine | 14.4268 | 3.50397 | 31.9039 |

Supplemental Figure 7. Specific productivity of amino acids of three respective EMS-treated mutants of *M. marburgensis* AHAS0 (A7, E3, E4) (n=2). Error bars mark the standard deviation.

**Definition of plating efficiency and survival rate of EMS-treated *M. marburgensis***

Supplemental Table 4: Plating efficiency *M. marburgensis*

| Plating efficiency [n=3] | CFU counts | CFU/mL | Counted cells | Plating efficiency |
| --- | --- | --- | --- | --- |
| 1:100 Dilution | lawn | n/a | 2.4·10^8^ | n/a |
| 1:10k Dilution | 50 +/- 3 | 5·10^7^ +/- 6% | 2.4·10^8^ | 20.87% |

CFU 2·105 cells

| EMS incubation / min | Colonies |
| --- | --- |
| 0 | 5000 |
| 7.5 | 2000 |
| 15 | 70 |
| 30 | 35 |
| 45 | 5 |
| 60 | 0 |

Supplemental Figure 8. Kill rate 1 mol L^-1^ EMS treatment with different incubation times.

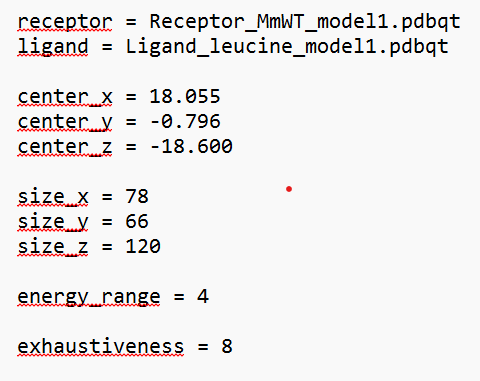

Supplemental Figure 9. Configuration sheet for Autodoc Vina docking assays of leucine in the IPMS-encoding genes of *E. coli*, *C. glutamicum*, and *M. marburgensis/M. thermautotrophicus*.

|  |  |
| --- | --- |
| methane | 91.3379% |
| single cell protein | 3.36468% |
| amino acids | 3.53836% |
| other organic compounds | 1.01146% |
| Intermediated and org acids | 0.747594% |

Supplemental Figure 10. Carbon balance of the fed-batch bioreactor fermentation with 100 L pilot scale. Percentages correspond to the amount of C-mol carbon in the outflow of the bioreactor broth and gas.

|  |  |  |
| --- | --- | --- |
| methane | 92.182% | 92.182% |
| single cell protein | 1.51853% | 1.51853% |
| amino acids | 3.47657% | 3.47657% |
| other organic compounds | 2.82286% | 2.82286% |

Supplemental Figure 11. Carbon balance of the fed-batch bioreactor fermentation with 10 L working volume. Percentages correspond to the amount of C-mol carbon in the outflow of the bioreactor broth and gas.
